# Genomic Surveillance of *Pseudomonas aeruginosa* in the Philippines from 2013-2014

**DOI:** 10.1101/2020.03.19.998229

**Authors:** Jeremiah Chilam, Silvia Argimón, Marilyn T. Limas, Melissa L. Masim, June M. Gayeta, Marietta L. Lagrada, Agnettah M. Olorosa, Victoria Cohen, Lara T. Hernandez, Benjamin Jeffrey, Khalil Abudahab, Charmian M. Hufano, Sonia B. Sia, Matthew T.G. Holden, John Stelling, David M. Aanensen, Celia C. Carlos, on behalf of the Philippines Antimicrobial Resistance Surveillance Program

## Abstract

*Pseudomonas aeruginosa* is an opportunistic pathogen often causing nosocomial infections that are resilient to treatment due to an extensive repertoire of intrinsic and acquired resistance mechanisms. In recent years, increasing resistance rates to antibiotics such as carbapenems and extended-spectrum cephalosporins have been reported, as well as multi-drug resistant and possible extremely drug-resistant rates of approximately 21% and 15%, respectively. However, the molecular epidemiology and AMR mechanisms of this pathogen remains largely uncharacterized.

We sequenced the whole genomes of 176 *P. aeruginosa* isolates collected in 2013-2014 by the Antimicrobial Resistance Surveillance Program. The multi-locus sequence type, presence of antimicrobial resistance (AMR) determinants, and relatedness between the isolates were derived from the sequence data. The concordance between phenotypic and genotypic resistance was also determined.

Carbapenem resistance was associated namely with loss-of function of the OprD porin, and acquisition of the metallo-β-lactamase VIM. The concordance between phenotypic and genotypic resistance was 93.27% overall for 6 antibiotics in 3 classes, but varied widely between aminoglycosides. The population of *P. aeruginosa* in the Philippines was diverse, with clonal expansions of XDR genomes belonging to multi-locus sequence types ST235, ST244, ST309, and ST773. We found evidence of persistence or reintroduction of the predominant clone ST235 in one hospital, as well as transfer between hospitals. Most of the ST235 genomes formed a distinct Philippine lineage when contextualized with international genomes, thus raising the possibility that this is a lineage unique to the Philippines. This was further supported by long-read sequencing of one representative XDR isolate, which revealed the presence of an integron carrying multiple resistance genes, including *bla*_VIM-2_, with differences in gene composition and synteny to other *P. aeruginosa* class 1 integrons described before.

We produced the first comprehensive genomic survey of *P. aeruginosa* in the Philippines, which bridges the gap in genomic data from the Western Pacific region and will constitute the genetic background to contextualize ongoing prospective surveillance. Our results also highlight the importance of infection control interventions aimed to curtail the spread of international epidemic clone ST235 within the country.

## Introduction

*Pseudomonas aeruginosa* is an opportunistic pathogen often causing nosocomial infections (e.g., pneumonia, bacteraemia and urinary tract infections), particularly in immunocompromised patients. ^1^ Eight Asian countries reported frequencies of isolation of *Pseudomonas* spp. above 15% from hospital-acquired pneumonia cases, with the Philippines reporting *P. aeruginosa* as the most common etiological agent. ^2^ In addition, *Pseudomonas* spp. were the second most common pathogen isolated from device-associated hospital acquired-infections in a study conducted across 9 intensive care units in three Philippine hospitals. ^3^

*P. aeruginosa* infections are resilient to treatment due to an extensive repertoire of intrinsic and acquired resistance mechanisms. ^4^ A strong association between the use of carbapenems and *P. aeruginosa* resistance has been documented. ^1^ Nonetheless, a recent study evaluating carbapenem restriction practices at a hospital in Manila found that 37% of the carbapenems prescriptions were non-compliant, thus pointing to ongoing challenges in antimicrobial stewardship. ^5^ Between 2010 and 2014, the Philippine Antimicrobial Resistance Surveillance Program (ARSP) reported increasing resistance rates to antibiotics used to treat *P. aeruginosa* infections such as carbapenems and extended-spectrum cephalosporins (Figure 1A-B). In contrast, resistance to aminoglycosides and fluoroquinolones has remained relatively stable or slightly decreased in the same period (Figure 1C). Alarmingly, the ARSP has also reported multi-drug resistant (MDR) rates of between 21 and 23%, and possible extremely drug-resistant (XDR) rates of between 13 and 18% in recent years. ^6–8^

**Figure 1.**
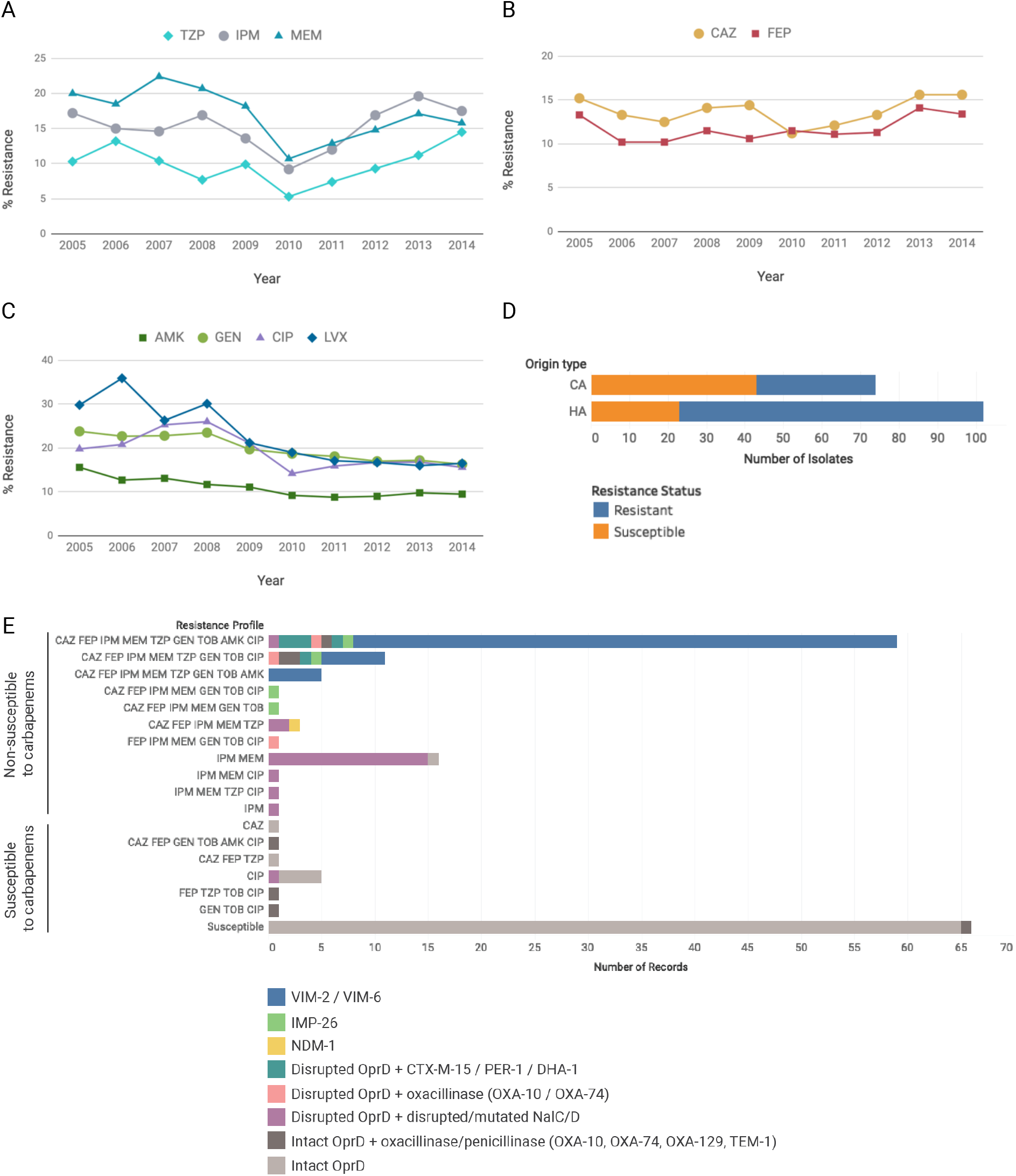
**A-C)** Yearly resistance rates to 9 antibiotics of *P. aeruginosa* isolates referred to ARSP 2005-2014. TZP: penicillin; IPM: imipenem; MEM: meropenem; CAZ: ceftazidime; FEP: cefepime; AMK: amikacin; GEN: gentamicin; CIP: ciprofloxacin; LVX: levofloxacin. **D)** Association between resistance and the origin of infection for 176 *P. aeruginosa* isolates sequenced in this study. CA: community-acquired; HA: hospital-acquired; Resistant to at least one antibiotic tested; Susceptible to all 9 antibiotics tested. **E)** Mechanisms of resistance to carbapenems and other beta-lactam antibiotics identified in the genomes of 176 isolates grouped by their resistance profile. For simplicity, only the main mechanism is indicated.

The emergence MDR *P. aeruginosa* that are resistant to carbapenems, aminoglycosides and fluoroquinolones was followed by reports of colistin-only sensitive isolates ^9^ and, more recently, of colistin resistance in carbapenem non-susceptible isolates. ^10^ All have a major impact on medical practice due to the few remaining treatment options. Interestingly, these reports coincide with multi-locus sequence type (ST) 235, ^9–11^ the predominant global epidemic clone. Several metallo-β-lactamase (MBL) genes have been identified in ST235 *P. aeruginosa* isolates from Asian countries, namely VIM and IMP, and usually associated with integrons carrying multiple resistance determinants. ^12–14^

While the resistance rates and profiles of *P. aeruginosa* in the Philippines have been characterized in detail by the ARSP and others, ^15, 16^ the molecular epidemiology and AMR mechanisms of this pathogen remained largely unknown. Whole-genome sequencing (WGS) can uncover transmission patterns, identify known AMR mechanisms, and pinpoint the source of acquired infections in the hospital setting. ^17^ In this study we characterized the clonal relatedness and resistance determinants of *P. aeruginosa* isolates collected by the ARSP between 2013 and 2014 using WGS.

## Methods

### Bacterial Isolates

A total of 7,877 *P. aeruginosa* isolates were collected and tested for resistance by the Philippine Department of Health - Antimicrobial Resistance Surveillance Program (DOH-ARSP) during the period of January 2013 to December 2014. Out of the 443 and 283 isolates referred to the Antimicrobial Resistance Surveillance Reference Laboratory (ARSRL) for confirmation in 2013 and 2014, respectively, 179 isolates from 17 sentinel sites were selected for whole-genome sequencing (WGS), as described in detail previously. ^18^ Additionally, we included 66 isolates susceptible to all antibiotics tested.

### Antimicrobial Susceptibility Testing (AST)

All *P. aeruginosa* isolates from this study were tested for antimicrobial susceptibility to 9 antibiotics representing 5 different classes, namely ceftazidime (CAZ), (FEP), imipenem (IPM), meropenem (MEM), piperacillin-tazobactam (TZP), gentamicin (GEN), tobramycin (TOB), amikacin (AMK) and ciprofloxacin (CIP) (Table 1). Antimicrobial susceptibility of the isolates was determined at the ARSRL using the Kirby-Bauer disk diffusion method, gradient methods such as E-Test (BioMerieux), and/or Vitek 2 Compact automated system (BioMerieux). The zone of inhibition and minimum inhibitory concentration of antibiotics were interpreted according to the 26^th^ edition of the Clinical and Laboratory Standard Institute (CLSI) guidelines to determine the resistance profile of the isolates as the list of antimicrobials to which the organism is non-susceptible. Multi-drug resistant phenotypes as classified according to recommended standard definitions. ^19^

**Table 1.** Total number of *P. aeruginosa* isolates analysed by the ARSP and referred to the ARSRL during 2013 and 2014, isolates submitted for whole-genome sequencing, and high-quality *P. aeruginosa* genomes obtained, discriminated by sentinel site and AMR profile.

|  | Number of Isolates |  |  |
| --- | --- | --- | --- |
|  | 2013 | 2014 | Total |
| **Total ARSP** | 3591 | 4286 | 7877 |
| **Referred to ARSRL** | 443 | 283 | 726 |
| **Submitted for WGS** | 89 | 90 | 179 |
| **High-quality genomes** | 87 | 89 | 176 |
| *By sentinel site* |  |  |  |
| BGH | 2 | 4 | 6 |
| BRH |  | 5 | 5 |
| CMC |  | 1 | 1 |
| CVM | 2 | 3 | 5 |
| DMC | 5 | 2 | 7 |
| EVR | 2 | 2 | 4 |
| FEU | 2 | 2 | 4 |
| GMH | 4 | 4 | 8 |
| JLM | 2 | 5 | 7 |
| MMH | 3 | 5 | 8 |
| NKI | 10 | 16 | 26 |
| NMC | 3 | 8 | 11 |
| RMC | 2 |  | 2 |
| SLH |  | 1 | 1 |
| STU | 5 | 4 | 9 |
| VSM | 32 | 16 | 48 |
| *By AMR profile* |  |  |  |
| Susceptible | 36 | 30 | 66 |
| CAZ FEP IPM MEM TZP GEN TOB AMK CIP [XDR] | 30 | 29 | 59 |
| IPM MEM | 7 | 9 | 16 |
| CAZ FEP IPM MEM TZP GEN TOB CIP [XDR] | 4 | 7 | 11 |
| CAZ FEP IPM MEM TZP GEN TOB AMK | 1 | 4 | 5 |
| CIP | 3 | 2 | 5 |
| CAZ FEP IPM MEM TZP | 1 | 2 | 3 |
| IPM MEM TZP CIP |  | 1 | 1 |
| GEN TOB CIP | 1 |  | 1 |
| FEP TZP TOB CIP |  | 1 | 1 |
| CAZ FEP IPM MEM GEN TOB | 1 |  | 1 |
| IPM | 1 |  | 1 |
| CAZ FEP IPM MEM GEN TOB CIP | 1 |  | 1 |
| IPM MEM CIP | 1 |  | 1 |
| CAZ FEP GEN TOB AMK CIP |  | 1 | 1 |
| FEP IPM MEM GEN TOB CIP |  | 1 | 1 |
| CAZ |  | 1 | 1 |
| CAZ FEP TZP |  | 1 | 1 |

### DNA Extraction and Whole Genome Sequencing

A total of 179 *P. aeruginosa* isolates were shipped to the Wellcome Trust Sanger Institute for whole-genome sequencing. DNA was extracted from a single colony of each isolate with the QIAamp 96 DNA QIAcube HT kit and a QIAcube HT (Qiagen; Hilden, Germany). DNA extracts were multiplexed and sequenced on the Illumina HiSeq platform (Illumina, CA, USA) with 100-bp paired-end reads. Isolate 13ARS-VSM740 was also sequenced with the PacBio RSII platform (Pacific Biosciences). Raw sequence data were deposited in the European Nucleotide Archive (ENA) under the study accession PRJEB17615. Run accessions for Illumina data are provided on the Microreact projects. The PacBio data was deposited under run accession ERR3284501.

### Bioinformatics Analysis

Genome quality was evaluated based on metrics generated from assemblies, annotation files, and the alignment of the isolates to the reference genome of *P. aeruginosa* strain LESB58 (accession FM209186), as previously described. ^18^ Assemblies were produced from short-read Illumina data as previously described, ^20^ and from long-read PacBio data with the HGAP v4 pipeline (Pacific Biosciences). A total of 176 isolates yielded high-quality *P. aeruginosa* genomes and were included in this study.

We derived *in silico* the multi-locus sequence type (MLST) of the isolates from the whole genome sequences. The sequence types (ST) were determined from assemblies with Pathogenwatch (https://pathogen.watch/) or from sequence reads with ARIBA ^21^ and the *P. aeruginosa* database hosted at PubMLST. ^22^ Integrons were detected in the genome assemblies with IntegronFinder. ^23^

Evolutionary relationships between isolates were inferred from single-nucleotide polymorphisms (SNPs) by mapping the paired-end reads to the reference genomes of *P. aeruginosa* strains LESB58 (FM209186) or NCGM2_S1(AP012280.1), as described in detail previously ^18^. Mobile genetic elements (MGEs) were masked from the alignment of pseudogenomes. For the phylogeny of ST235 genomes, recombination regions detected with Gubbins ^24^ were also removed. Alignments of SNPs were inferred with snp-sites v2.4.1 ^25^, and were also used to compute pairwise SNP differences between isolates from different patients (minimum N=3) belonging to the same or to different hospitals with an in-house script. Maximum likelihood phylogenetic trees were generated with RAxML ^26^ based on the generalised time reversible (GTR) model with GAMMA method of correction for among-site rate variation and 100 bootstrap replications. The tree of 904 global *P. aeruginosa* genomes with geolocation and isolation date for which raw sequence data are available at the European Nucleotide Archive was inferred using an approximately-maximum likelihood phylogenetic method with FastTree. ^27^

Known AMR determinants were identified from raw sequence reads using ARIBA ^21^ and a curated database of known resistance genes and mutations ^28^. Results were complemented with predictions using the Comprehensive Antibiotic Resistance Database (CARD ^29^), and a custom database of mutations in the quinolone resistance-determining region of the *gyrA/B* and *parC/E* genes described for *P. aeruginosa* ^4^. Genomic predictions of resistance were derived from the presence of known antimicrobial resistance genes and mutations identified in the genome sequences. The ARIBA output for the porin gene *oprD* was inspected to detect loss-of-function mutations (frameshifts, premature STOP codons, failed START or STOP codons). The *oprD* sequences were extracted from the whole-genome draft assemblies with blastn using the *oprD* sequence from strain PAO1 (accession NC_002516.2, genome positions 1043982-1045314) as a query. The *oprD* DNA sequences thus obtained were translated *in silico* to inspect the integrity of the coding frames. A 444 or 442 amino-acid protein including a START and a STOP codon was considered functional.

The genomic predictions of AMR (test) were compared to the phenotypic results (reference) and the concordance between the two methods was computed for each of 6 antibiotics (1056 total comparisons). Isolates with either a resistant or an intermediate phenotype were considered resistant for comparison purposes.

All project data, including inferred phylogenies, AMR predictions and metadata were made available through the web application Microreact (http://microreact.org).

## Results

### Demographic and Clinical Characteristics of the *P. aeruginosa* Isolates

Out of the 179 *P. aeruginosa* isolates submitted for WGS, 176 passed quality control filters and were confirmed as *P. aeruginosa in silico*. The demographic and clinical characteristics of the 176 *P. aeruginosa* isolates are summarized in Table 2. The age range of the patients were < 1 up to 96 years old, with 27% (N=47) of the isolates from patients 65 years old and above. Fifty-eight percent (N=102) of the isolates were classified as hospital-acquired infection isolates. As to specimen type, 53% (N=94) of isolates were isolated from respiratory samples (tracheal aspirates and sputum), followed by wound (N=25), blood (N=21), and urine (N=12).

**Table 2.**
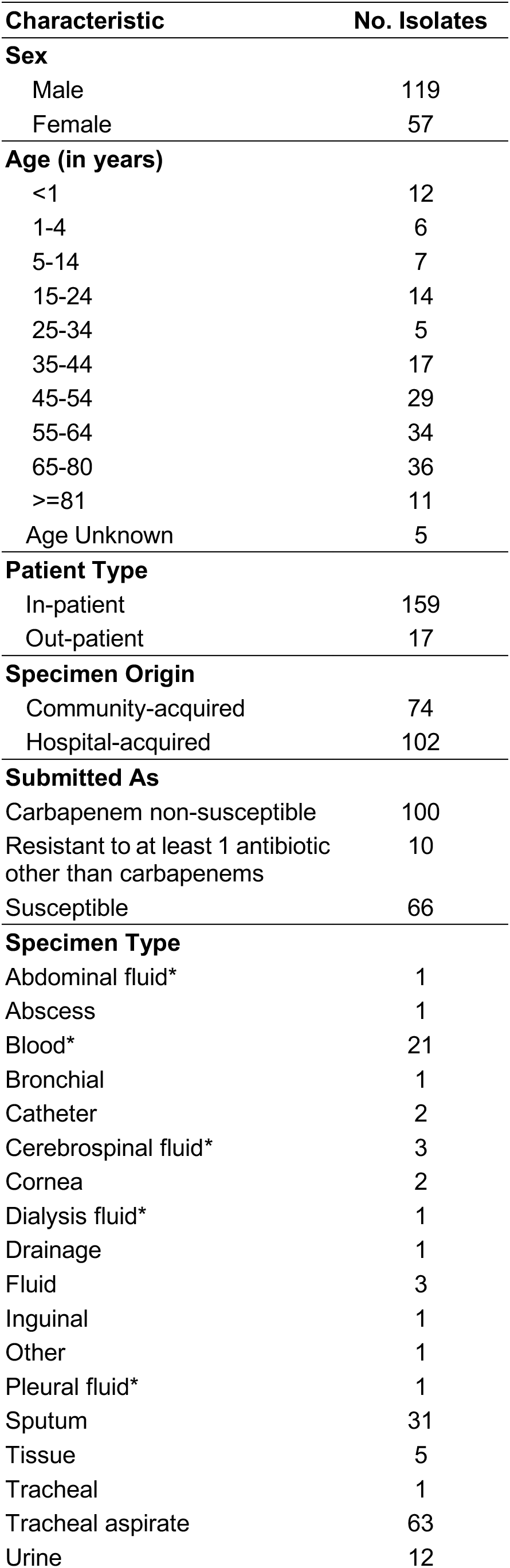
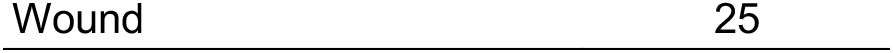
Demographic and clinical characteristics of 176 *P. aeruginosa* isolates. Invasive isolates were considered as those obtained from specimen types marked with an asterisk (*).

| **Characteristic** | **No. Isolates** |
| --- | --- |
| **Sex** |  |
| Male | 119 |
| Female | 57 |
| **Age (in years)** |  |
| <1 | 12 |
| 1-4 | 6 |
| 5-14 | 7 |
| 15-24 | 14 |
| 25-34 | 5 |
| 35-44 | 17 |
| 45-54 | 29 |
| 55-64 | 34 |
| 65-80 | 36 |
| >=81 | 11 |
| Age Unknown | 5 |
| **Patient Type** |  |
| In-patient | 159 |
| Out-patient | 17 |
| **Specimen Origin** |  |
| Community-acquired | 74 |
| Hospital-acquired | 102 |
| **Submitted As** |  |
| Carbapenem non-susceptible | 100 |
| Resistant to at least 1 antibiotic other than carbapenems | 10 |
| Susceptible | 66 |
| **Specimen Type** |  |
| Abdominal fluid* | 1 |
| Abscess | 1 |
| Blood* | 21 |
| Bronchial | 1 |
| Catheter | 2 |
| Cerebrospinal fluid* | 3 |
| Cornea | 2 |
| Dialysis fluid* | 1 |
| Drainage | 1 |
| Fluid | 3 |
| Inguinal | 1 |
| Other | 1 |
| Pleural fluid* | 1 |
| Sputum | 31 |
| Tissue | 5 |
| Tracheal | 1 |
| Tracheal aspirate | 63 |
| Urine | 12 |

### Concordance Between Phenotypic and Genotypic AMR

Isolates were tested for susceptibility to 9 antibiotics representing 5 different classes (Figure 1A-C, and Table 3). As for other organisms in the See and Sequence study, our sequencing survey was focused on isolates non-susceptible to carbapenems (N=100). Only 10 isolates susceptible to carbapenems but resistant to other antibiotics were included (Table 1). However, unlike for other organisms, 66 *P. aeruginosa* isolates susceptible to all 9 antibiotics tested were available and were also included in the sequencing survey. We observed a significant association between the resistance status of the isolates (susceptible or resistant) and their classification according to origin of infection (community- or hospital-acquired, two-tailed Fisher’s exact test *p*=0.000002). Community-acquired infections were more frequently associated with susceptible isolates, while hospital-acquired infections with resistant isolates (Figure 1D).

**Table 3.** Comparison of genomic predictions of antibiotic resistance with laboratory susceptibility testing at the ARSRL.

| Antibiotic Class | Antibiotic | Isolates Tested | Resistant Isolates (AST) | False Positive | False Negative | % Concordance | Acquired Resistance Mechanisms |
| --- | --- | --- | --- | --- | --- | --- | --- |
| Carbapenem | Imipenem | 176 | 100 | 1 | 4 | 97.16 | *bla*<sub>VIM-2</sub>, *bla*<sub>VIM-6</sub>, *bla*<sub>NDM-1</sub>, *bla*<sub>IMP-26</sub>, OprD loss-of-function (*oprD* interrupted, fragmented, or missing, presence of premature STOP, START codon missing), NalC/D loss-of-function (*nalC* missing, NalC_G71E, S209R, A186T, NalD_S32N) |
|  | Meropenem | 176 | 99 | 2 | 4 | 96.59 |  |
| Aminoglycoside | Gentamicin | 176 | 77 | 0 | 34 | 80.68 | *aac(3)-II*, *aac(6')-31*, *aac(6')-IIa*, *ant(2'')-Ia* |
|  | Tobramycin | 176 | 78 | 2 | 3 | 97.16 | *aac(3)-II*, *aac(6')-31*, *aac(6')-Ib*, *aac(6')-Ib-cr*, *aac(6')-IIa*, *ant(2'')-Ia* |
|  | Amikacin | 176 | 61 | 14 | 4 | 89.77 | *aac(6')-31*, *aac(6')-Ib*, *aac(6')-IIa*, *aph(3')-VI* |
| Fluoroquinolone | Ciprofloxacin | 176 | 82 | 5 | 12 | 93.75 | *qnrD*, *qnrVC*, *aac(6')-Ib-cr*, GyrA_D87N, D87Y, T83I, GyrB_E468D, S466F, ParC_S87L |

Seventeen out of 18 isolates resistant to imipenem and meropenem but not to other beta-lactam antibiotics carried both loss-of-function disruptions in the OprD porin gene, and disruptions or known non-synonymous mutations in the NalC (A186T, G72E, S209R) and/or NalD (S32N) regulators of the MexAB-OprM multi-drug efflux pump, suggesting that their resistance is owed to a combination of reduced influx and increased efflux of the carbapenem antibiotics, respectively (Figure 1E). From the 81 carbapenem-resistant isolates that were also resistant to third-generation cephalosporin ceftazidime and/or fourth-generation cephalosporin cefepime, 67 isolates carried acquired metallo-β-lactamase (MBL) genes *bla*_VIM-2_ (N=61 genomes), *bla*_VIM-6_ (N=1), *bla*_IMP-26_ (N=4), or *bla*_NDM-1_ (N=1), while 5 carried disrupted *oprD* genes plus acquired extended-spectrum β-lactamase (ESBL) genes *bla*_PER-1_ (N=3), *bla*_CTX-M-15_ (N=1) or AmpC-like gene *bla*_DHA-1_ (N=1). The remaining 8 isolates harboured other β-lactamase genes but their carbapenem resistance mechanisms remain uncharacterized. Seventy-five of the 76 isolates susceptible to carbapenems carried either a full-length OprD porin (444 amino acids) without any known mutations, or a 442 amino acid-long OprD protein with an intact reading frame, while one isolate was missing the STOP codon in the *oprD* gene.

Comparisons between phenotypic and genotypic data for are presented for carbapenems, aminoglycosides and fluoroquinolones in Table 3. The overall concordance was 93.27% for the 6 antibiotics analysed, with the lowest concordance exhibited by gentamicin (80.68%) and the highest for tobramycin (97.16%). The concordance was above 96% for carbapenems.

### Genotypic Findings

#### *In silico* genotyping

Multi-locus sequence type was predicted *in silico* from the whole-genome sequence data of the 176 *P. aeruginosa* isolates. A total of 79 STs were identified, with 27.8% (N=49) of the isolates belonging to ST235, followed by ST309 (5.7%, N=10), ST244, and ST773 (5.1% each, N=9). The majority of the STs (79.7%, N=63) were singletons (represented by only one genome), most of which (N=42) were contributed by the susceptible isolates. Indeed, the clonal diversity of the resistant isolates (36 STs, N=110) was lower than that of the susceptible isolates (56 STs, N=66). ST235 represented 43.6% (N=48) of the resistance isolates but only 1.5% (N=1) of the susceptible isolates, and was predominantly a nosocomial clone in the Philippines (36 HAI vs 13 CAI isolates), spread geographically across 13 hospitals. Details of the number and most common sequence types (STs) found in each of the sentinel sites are shown on Table 4.

**Table 4.**
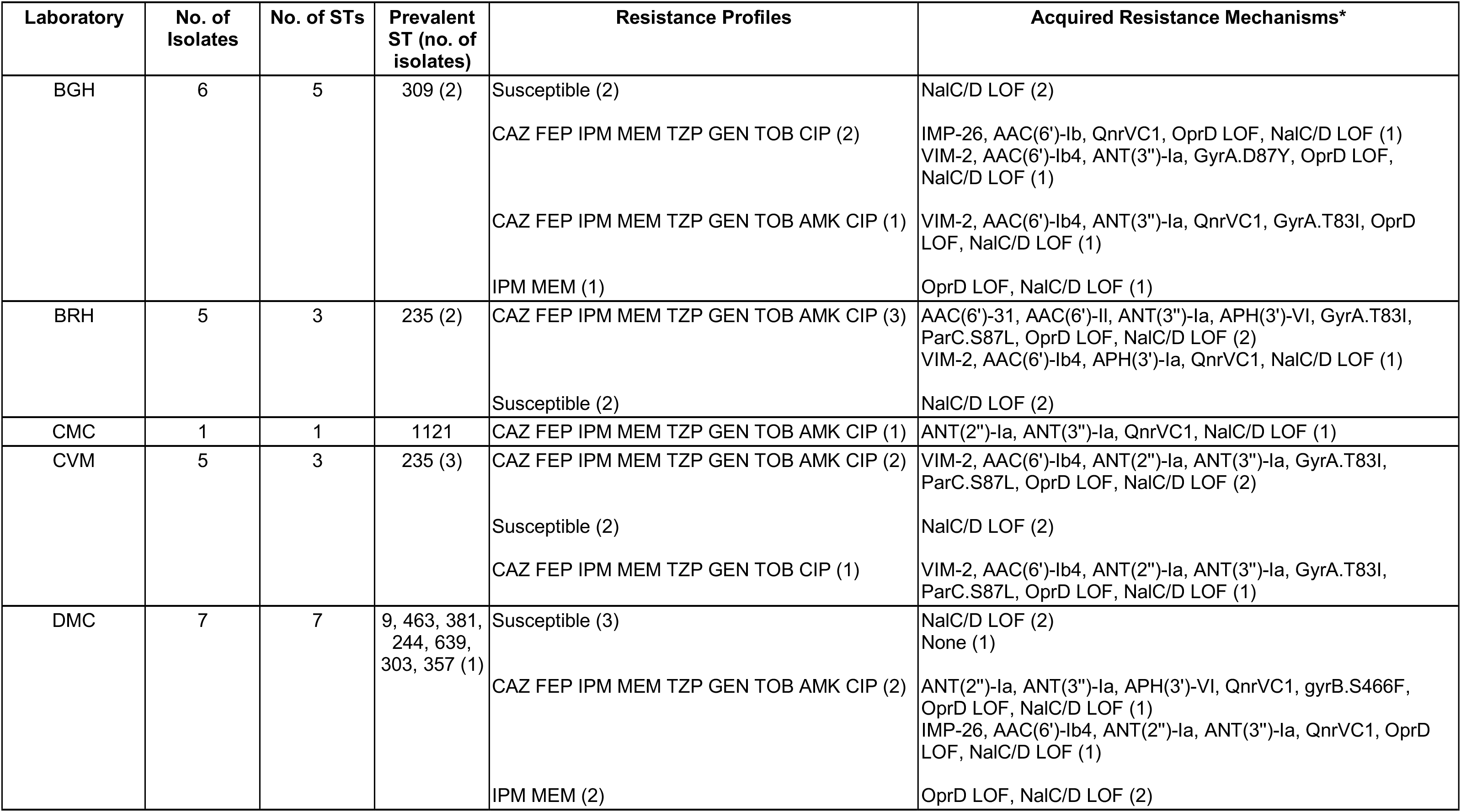
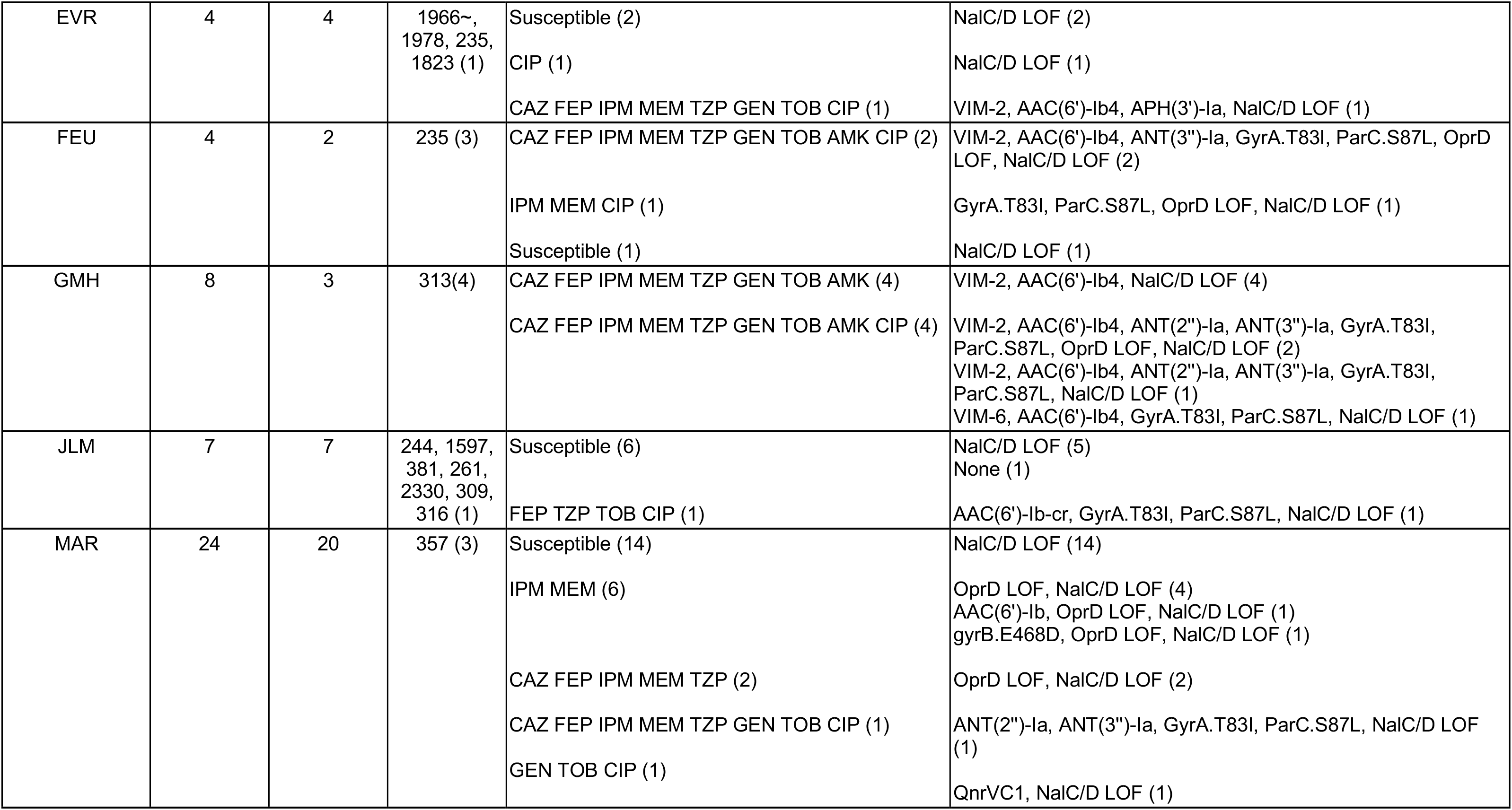
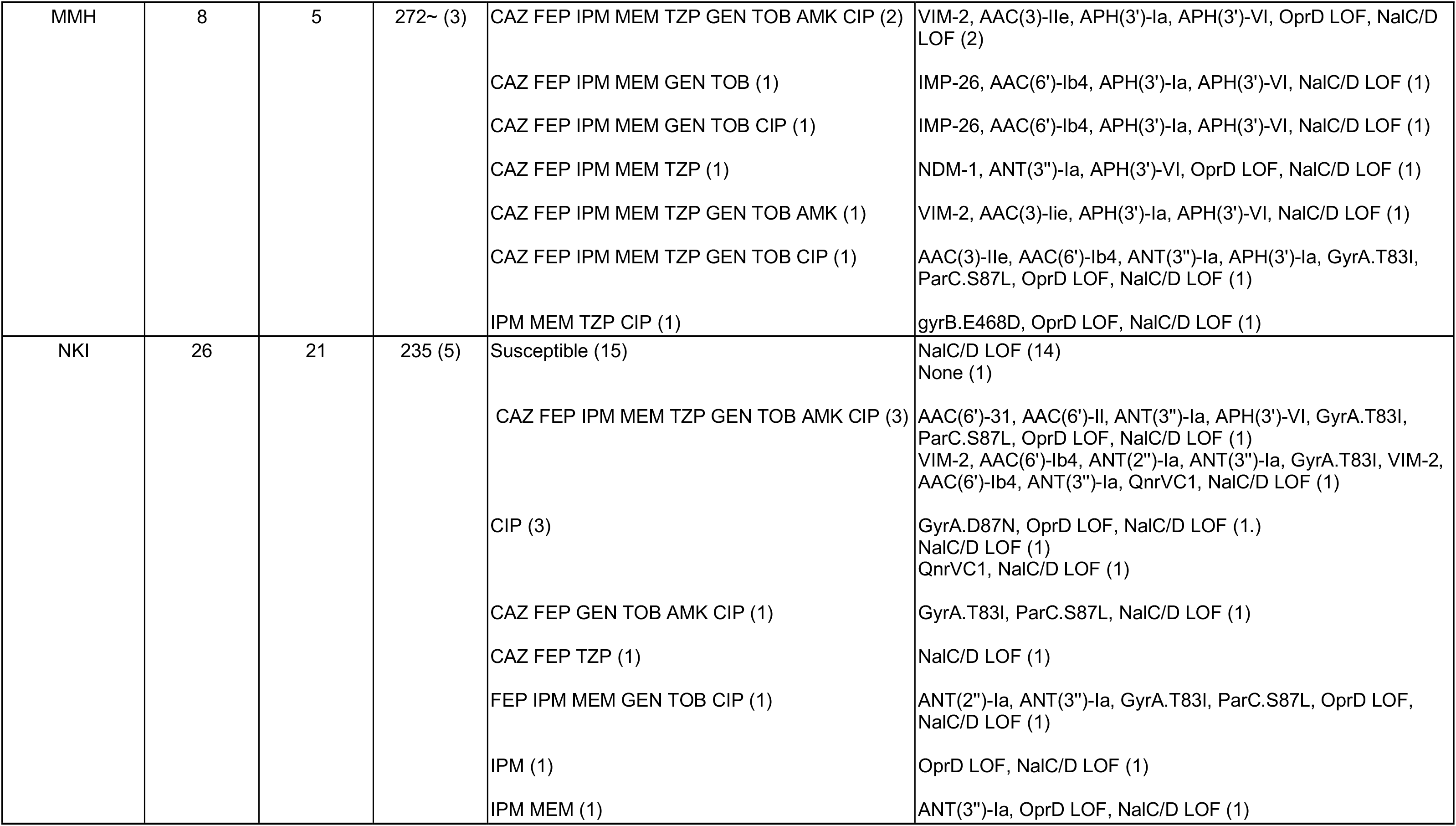
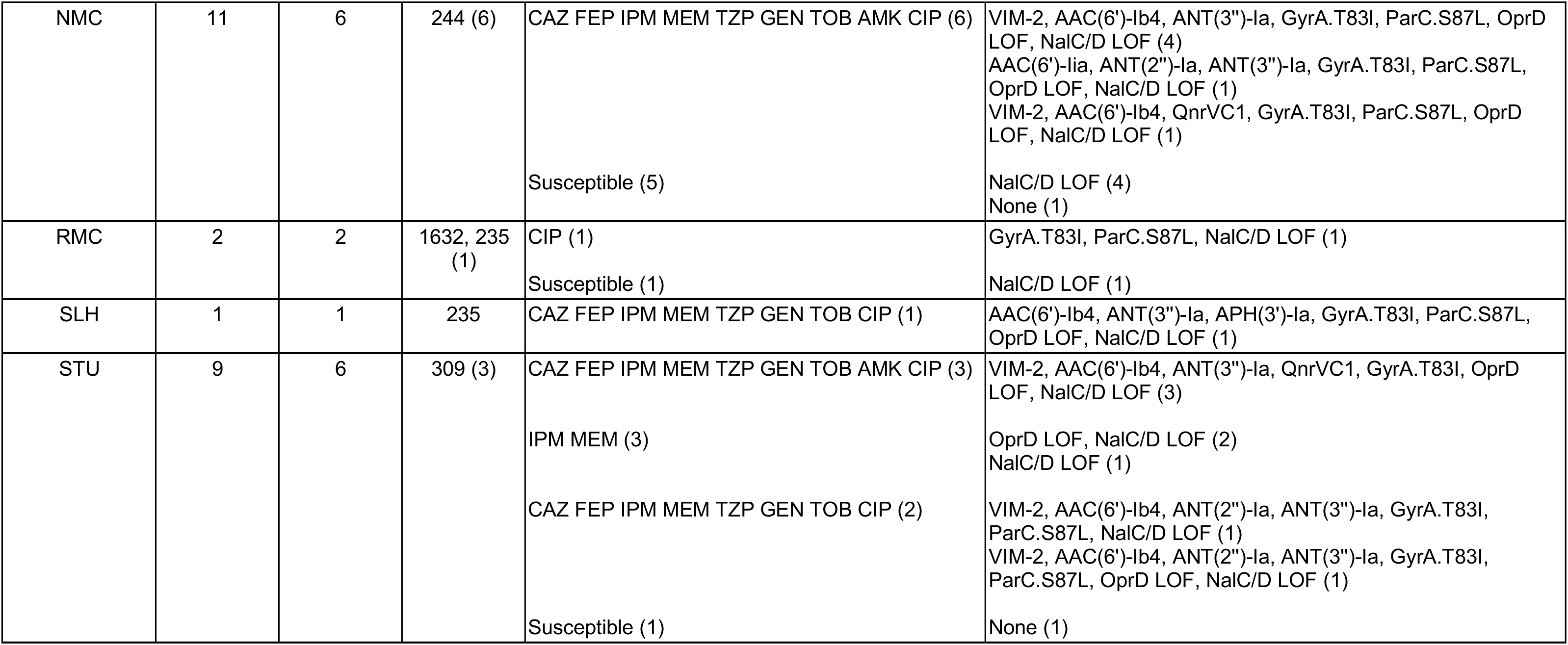
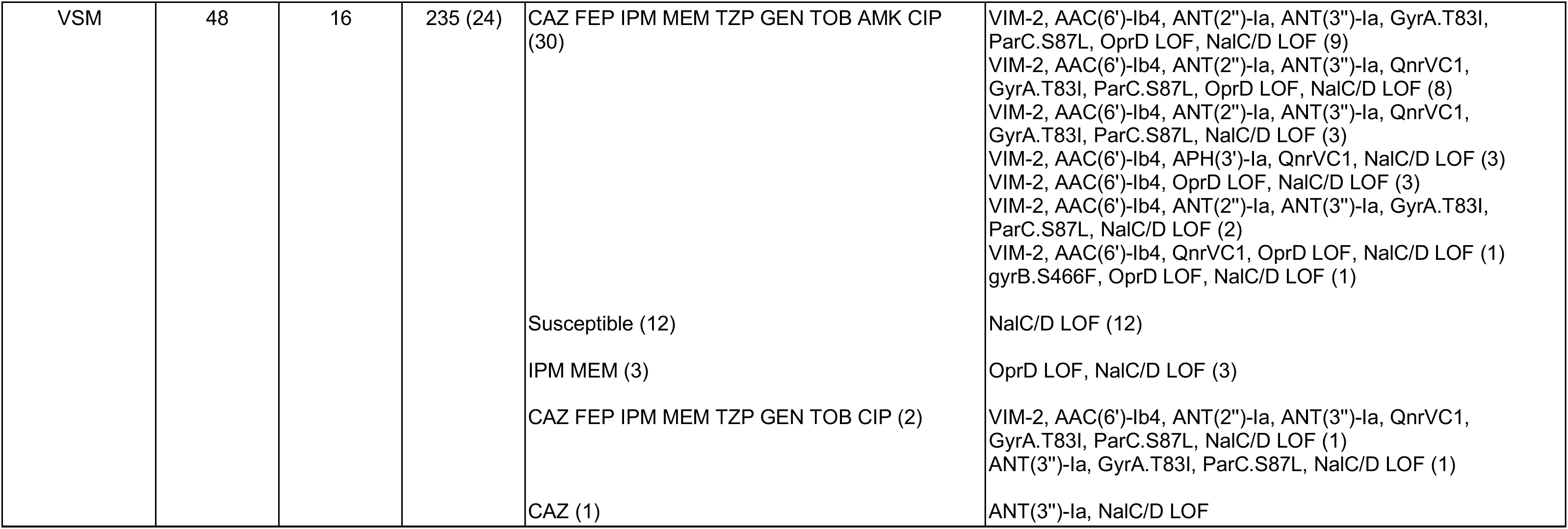
Distribution of isolates, sequence types (STs), resistance profiles and acquired resistance mechanisms across the 17 sentinel sites. Only genes and mutations associated with the antibiotic classes tested are shown (beta-lactamases, aminoglycosides, and fluoroquinolones). The full complement can be found in https://microreact.org/project/ARSP_176PAE_2013-2014. LOF: loss-of-function.

#### Population structure of *P. aeruginosa* in the Philippines

The phylogenetic tree of 176 genomes from the Philippines was composed of three major groups previously described (Freschi 2015), group 1 (N=64) including PA14, group 2 (N=105) including PAO1, and the more distantly related group 3 (N=7) including PA7 (Figure 2A). All three groups included carbapenem-resistant isolates and susceptible isolates, though the majority of isolates in group 2 were susceptible (N=39, 60.9%), and the majority of isolates in group 1 were carbapenem-resistant (N=75, 71.4%, Figure 2B).

**Figure 2.**
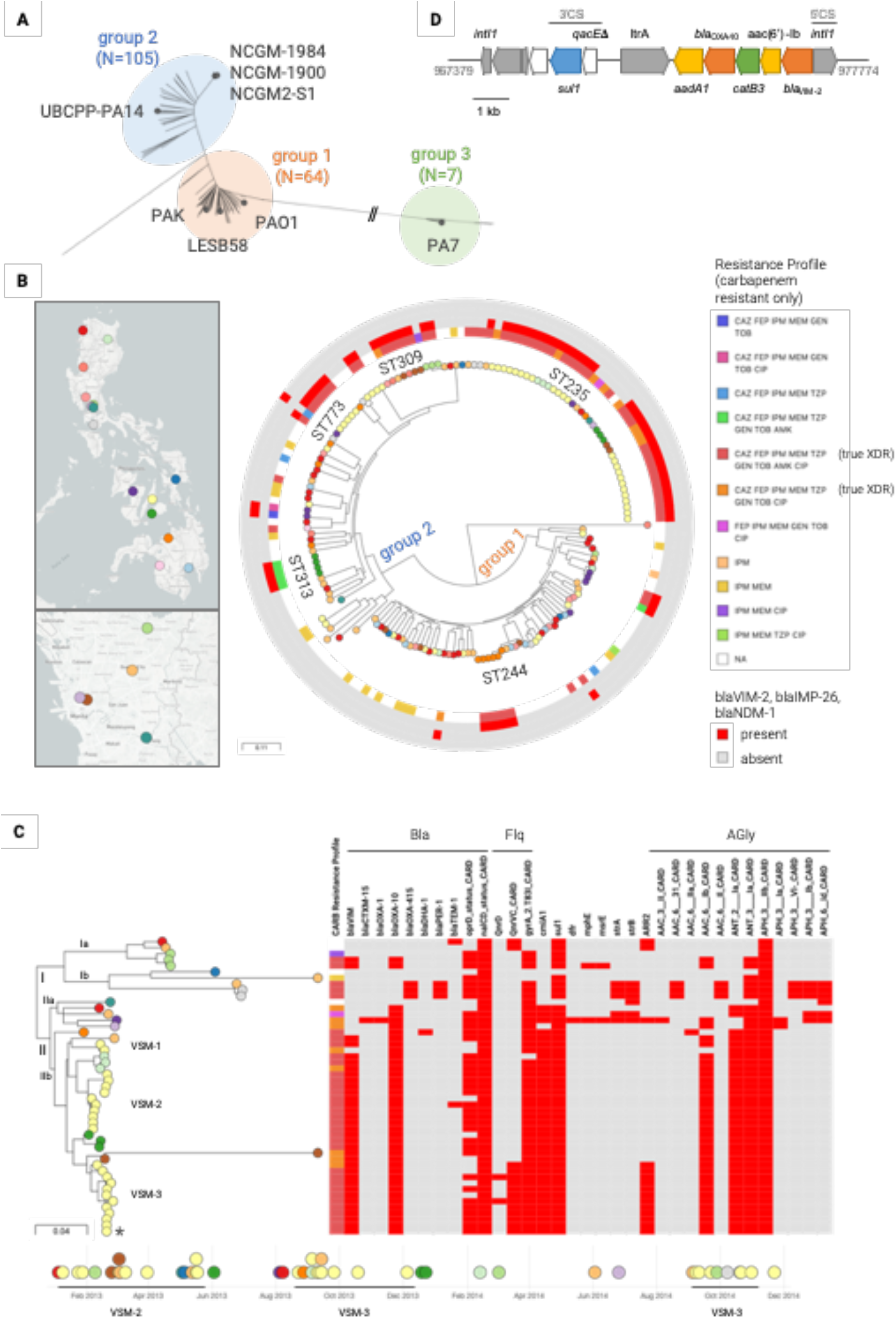
Genomic surveillance of *P. aeruginosa* from the Philippines 2013-2014. **A)** Phylogenetic tree of 176 isolates from the Philippines and 8 reference genomes, inferred with RAxML from an alignment of 396,194 core SNP sites. The reference genomes are indicated by grey nodes. **B)** Phylogenetic tree of 169 isolates from groups 1 and 2 inferred with RAxML from an alignment of 305,220 SNP sites obtained after mapping the genomes to the complete genome of strain LESB58 (ST146) and masking mobile genetic elements from the alignment. The tree leaves are coloured by sentinel site and indicated on the map (left panels, top: Philippines, bottom: detail of the National Capital Region). Inner ring: Sequence Type. Tree rings indicate (from inner to outer) the distribution of the carbapenem resistant profiles and of carbapenemase genes *bla*_VIM_, *bla*_IMP_ and *bla*_NDM_. The data, including the full distribution of resistance determinants, is available at https://microreact.org/project/ARSP_169PAE_2013-2014. **C)** Phylogenetic tree of 49 ST235 genomes inferred from an alignment of 1,066 SNP sites obtained after mapping the genomes to the complete genome of strain NCGM2-S1 (ST235) and masking mobile genetic elements and recombination regions. The tree leaves are coloured by sentinel site as indicated on the map from Figure 2B. The tree blocks represent the distribution of the carbapenem resistant profiles and of acquired resistance genes and mutations. Two closely related isolates isolated 1.5 years apart are indicated with arrowheads on the tree and on the timeline. The representative isolate sequenced with long reads is shown with an asterisk. The full data is available at https://microreact.org/project/ARSP_PAE_ST235_2013-14. The scale bars represent the number of single nucleotide polymorphisms per variable site. **D)** Resistance genes acquired en bloc within an class 1 integron in *P. aeruginosa* strain 14ARS-VSM0870. Arrows indicate genes conferring resistance to beta-lactamases (orange), aminoglycosides (yellow), chloramphenicol (green) and sulphonamides (blue), or related to DNA mobilization/integration (grey). 3’CS and 5’CS: conserved segments.

The population of *P. aeruginosa* is composed of a limited number of widespread clones that are selected from a diverse pool of rare, unrelated genotypes that recombine at high frequency. ^30^ A phylogenetic tree of 169 genomes from groups 1 and 2 showed that the clonal expansions were found mostly within the major group 1, and were represented by ST235, ST309, ST773, and ST313 (Figure 2B), which were found across multiple hospitals and presented resistance to multiple antibiotics. The majority of the XDR isolates (N=61, 87%) were found in ST235, ST244, ST309, and ST773, and most of them (N=44, 62.8%) carried *bla*_VIM_ (an MBL that can degrade all anti-pseudomonal beta-lactamases except for aztreonam),^1^ AAC(6’)-Ib (an aminoglycoside acetyltransferase conferring resistance to tobramycin and amikacin), and GyrA_T83I (a mutation associated with resistance to fluoroquinolones), among several other genes and mutations.

The higher prevalence of ST235 prompted us to look into this clone in more detail. The phylogenetic tree of 49 ST235 isolates showed that it was composed of two distinct clades with different geographic distribution (Figure 2C). Clade I (N=10) was represented in 5 hospitals in the Luzon (north) and Visayas (central) island groups, while clade II (N=39) was represented in 10 hospitals from north to south of the country. The phylogeographic structure of the tree and the relatedness between genomes showed evidence of dissemination of ST235 between hospitals. Within clade Ib (Figure 2C) we found one genome from NKI that differed from two genomes from BHR by 7 and 8 SNPs, respectively. Within clade IIb (Figure 2C) the pairwise SNP differences ranged between 0 and 64 for isolates from the same hospital (mean 36.41 ± 20.84), and between 29 and 61 for isolates from different hospitals (mean 45.36 ± 8.12). The lack of significant differences between the two groups (Mann-Whitney U test z-score=-1.49145, *p*=0.13622) implied that the clusters of genomes from different hospitals observed on the tree are closely related.

The genomes from the VSM hospital (N=24) formed at least three clusters within clade IIb, two of which exhibited discreet temporal distribution (VSM-2 and VSM-3, Figure 2C), suggesting that they could represent hospital outbreaks. For example, the genomes from different patients within clade VSM-3 differed by an average of 11.55 pairwise SNPs (range 0 to 24). We also identified isolates from different patients within VSM-3 that were collected between nine or more months apart (Figure 2C), suggesting that ST235 can either persist in or be reintroduced to the hospital environment.

The distribution of acquired resistance genes and mutations showed the presence of different complements of resistance determinants between clades I and II, with patterns that are consistent with the acquisition of multiple genes simultaneously by mobile genetic elements. Long-read sequencing of isolate 14ARS-VSM0870 representative of the XDR resistant profile CAZ FEP IPM MEM TZP GEN TOB AMK CIP (marked with an asterisk on Figure 2C), revealed the acquisition of *bla*_VIM-2_, *bla*_OXA-10_, *catB3*, *aadA1* (ANT(3”)-Ia) and *acc(6’)-Ib* within a class 1 integron integrated in the chromosome at position 977,774 (Figure 2D), while the ciprofloxacin resistance gene *qnrVC* and the rifampin-resistance gene *arr-2* were located on a different class I integron elsewhere in the genome.

#### *P. aeruginosa* from the Philippines in global context

We placed the genomes from our retrospective collection in the global context of 904 contemporary *P. aeruginosa* public genomes available from sequence data archives with linked geographic and temporal information, and collected mainly between 2007 and 2017. This collection of public genomes represents 17 countries and 178 STs, but more than 60% of the genomes were from Europe (N=373) and USA (N=205). The Philippine genomes were found throughout the tree, indicating that the *P. aeruginosa* population captured in our survey largely represents the global diversity of this pathogen, but with the notable absence of several high-risk clones. Sequence types ST235, ST111 and ST175 are the dominant international epidemic clones responsible for MDR and XDR nosocomial infections worldwide, and known carriers of MBL genes. However ST111 and ST175 was not found in our survey (Figure 3A). Other recognized high-risk clones, such as ST233 and ST773 carrying *bla*_VIM-2_ or *bla*_IMP-1_ have been reported from countries in Asia, Africa and Europe (Wright 2015). ST233 was represented by only one genome from the Philippines without any MBL genes and positioned on a separate, long branch within the tree, indicating a more distant relationship to the 66 ST233 isolates from other countries. ST773 was represented by nine genomes from the Philippines, carrying either *bla*_VIM-2_ (N=6), *bla*_NDM-1_ (N=1) or no MBL gene (N=2), but our collection did not include any ST773 genomes from other countries.

**Figure 3.**
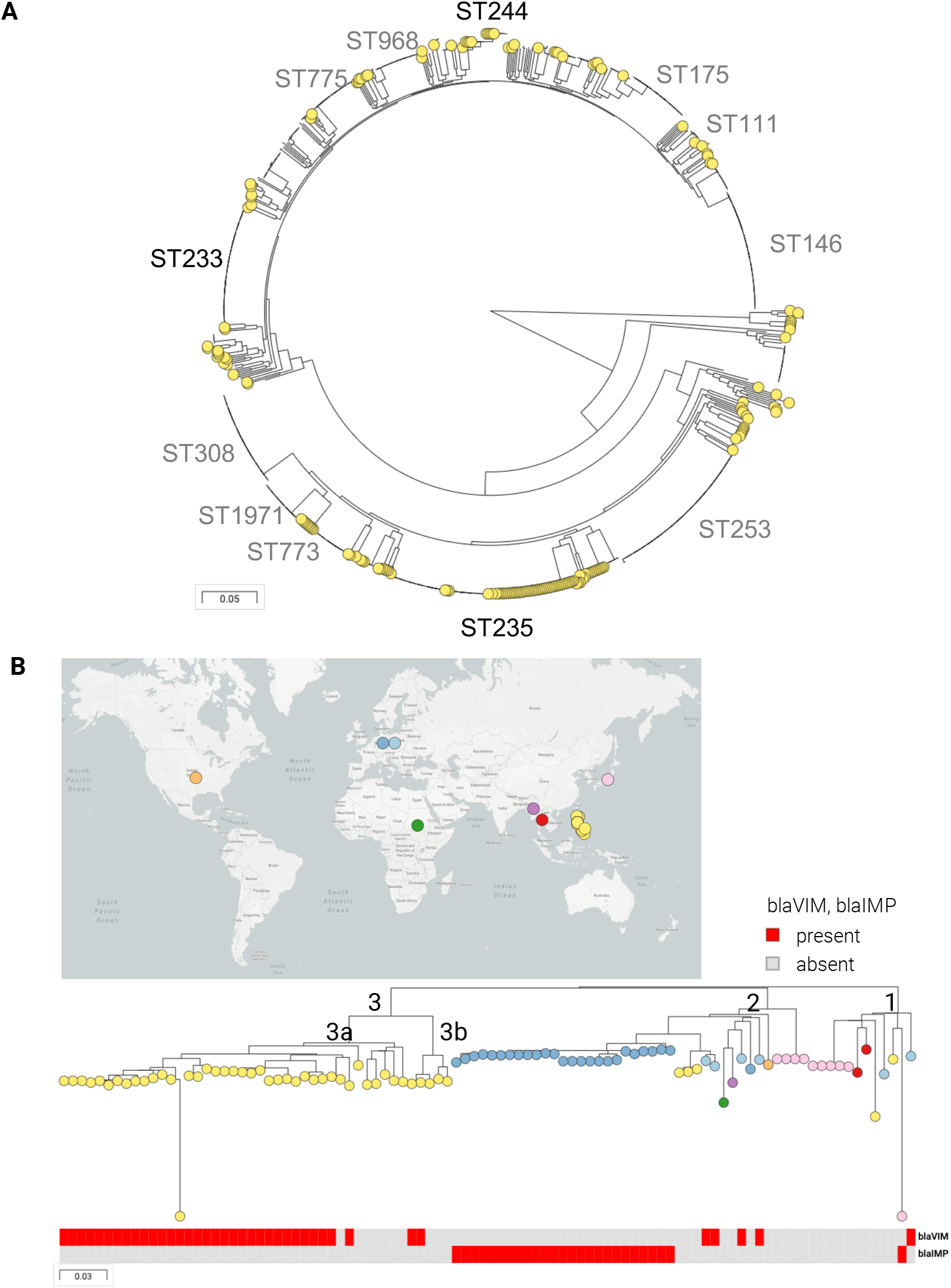
*P. aeruginosa* from the Philippines in global context. **A)** Phylogenetic tree of 904 *P. aeruginosa* isolates from the Philippines (N=176, this study) and from 57 other countries inferred from an alignment of 549,126 SNP positions obtained after mapping the genomes to the complete genome of strain LESB58 and masking regions with mobile genetic elements. The yellow tree nodes indicate the genomes from this study. The major lineages (STs) are labelled in black if represented by genomes of this study, or in grey if they are not. The data is available at https://microreact.org/project/ARSP_PAE_GLOBAL. **B)** Phylogenetic tree of 96 ST235 isolates inferred from an alignment of 1,993 SNP positions obtained after mapping the genomes to the complete genome of strain NCGM2-S1 (ST235) and masking mobile genetic elements and recombination regions. The tree leaves are coloured by country as indicated on the map. The tree blocks represent the distribution of acquired carbapenemase genes. The data is available at https://microreact.org/project/ARSP_PAE_ST235_GLOBAL. The scale bars represent the number of single nucleotide polymorphisms (SNPs) per variable site.

A more detailed tree of 96 ST235 genomes of global distribution showed three major clades, clade 1 represented by isolates from Germany, Japan, Thailand and the Philippines (N=2), clade 2 with the broadest geographic distribution across four continents and also including isolates from this study (N=3), and clade 3 composed exclusively of isolates from the Philippines (N=44, Figure 3B). The observation that the majority of the Philippine genomes (89.8%) formed a distinct cluster without any closely related genomes from other countries (Figure 3B) raises the possibility that this is a sub-lineage of ST235 characteristic to the Philippines. However, introductions from the other globally dispersed lineages may also occur, as shown in clades 1 and 2.

## Discussion

In this study we combined WGS and laboratory-based surveillance to characterize susceptible and resistant *P. aeruginosa* circulating in the Philippines in 2013 and 2014 with a particular emphasis on resistance to carbapenems, which we showed to be on the rise in the years preceding this survey (Figure 1A). Drug-resistant *P. aeruginosa* infections are more difficult to treat, resulting in poor patient outcomes. A previous study conducted in a tertiary hospital in Manila showed that severity of illness and mortality rates were significantly higher among patients infected with drug-resistant *P. aeruginosa* compared to susceptible isolates, while median duration of hospital stay was significantly longer. ^31^

*P. aeruginosa* strains exhibit a complex interplay between resistance mechanisms, both intrinsic and acquired. ^32^ The current uneven understanding of some of the resistance mechanisms was reflected in the variable concordance between phenotypic and genotypic resistance for the different antibiotics (Table 3), even for those antibiotics belonging to the same class (aminoglycosides). Our characterization of the carbapenem resistance showed a combination of diverse known mechanisms, from inhibition of antibiotic influx into the cell (OprD loss-of-function mutations), upregulation of antibiotic efflux out of the cell (NalC/D loss-of-function mutations), to carbapenemase genes, namely metallo-β-lactamase VIM and IMP (Tables 3 and 4). Although additional mechanisms may exist, the concordance between phenotypic and genotypic predictions of AMR was high for the carbapenems, but it required a degree of curation of results that is not practical within public health settings.

There are clear limitations in the genomic predictions of AMR for *P. aeruginosa.* Firstly, publicly-available, curated databases are not comprehensive of all the known mechanisms. For example, we complemented the prediction of genomic resistance to ciprofloxacin with a custom database of known mutations. ^4^ Similarly, no mutations leading to upregulation of the chromosomal cephalosporinase AmpC (*bla*_PAO_) were found in our data, but an exhaustive search would require additional analyses. Secondly, the regulatory pathways of some mechanisms are not fully understood, such as those that regulate AmpC. ^32, 33^ Thirdly, extensive manual curation of some of the predictions is needed to ensure accuracy, for example of the loss-of-function mutations in the *oprD* gene.

The most prevalent clone in our dataset was ST235 (27.8% of the isolates, N=49), found throughout the Philippines. ST235 is a well-characterized international epidemic clone causing drug-resistant nosocomial outbreaks. ^30^ Isolates carrying *bla*_VIM-2_ and belonging to ST235 were reported from Malaysia, Thailand, and Korea. ^13^ With the resolution of WGS, we showed evidence of potential localized hospital outbreaks of ST235, as well as of persistence/reintroduction of this clone within one hospital. The number of SNP differences between genomes of isolates from different patients (0-24) were consistent with those reported for a persistent outbreak of *P. aeruginosa* in a hospital the United Kingdom. ^34^ We also showed evidence of transfer of ST235 between hospitals (Figure 2C), with isolates from different hospitals separated by as little as 7 SNPs. Patient transfer between hospitals is not common in the Philippines, but the sampling of this study only allows us to hypothesize about a possible role of the community, animals and/or the environment in spread of this clone.

It was previously proposed that ST235 emerged in Europe around 1984, coinciding with the introduction of fluoroquinolones, and disseminated to other regions via two independent lineages, acquiring resistance determinants to aminoglycosides and beta-lactams locally. ^14^ Simultaneous acquisition of resistance to multiple antibiotics via integrons, transposons and integrative conjugative elements is well described in *P. aeruginosa*, ^35^ which was also apparent in the distribution of resistance genes in our genomes (Figure 2C). We have shown an example of a class I integron carrying six resistance genes in the genetic background of ST235 (Figures 2C-D). While this integron shared some features with others previously described in *P. aeruginosa*, ^13, 30^ such as the 5’ and 3’ conserved segments, ^35^ the gene composition and synteny was different, thus supporting the hypothesis of local acquisition of resistance.

Country-specific ST235 lineages have been reported before, ^11, 14^ confirming that country-wide clonal expansions may occur in the context of the global circulation of this clone. A previous longitudinal study also showed VIM-2 positive ST235 spreading throughout Russia, Belarus and Kazakhstan, albeit without the resolution of whole genome data. ^36^ The contextualization of our genomes with international ST235 genomes showed a distinct cluster of Philippine genomes, suggesting that it could be specific to the country (Figure 3B). Alternatively, this could be explained by the limited representation of the Western Pacific region in the collection of global genomes, which, in turn, points to the need for public genome data with more even geographical coverage. Our retrospective survey contributed to bridge this gap by making raw sequence data readily available on public archives.

In conclusion, our detailed description of the epidemiology and resistance mechanisms of ST235 in the Philippines (the first to our knowledge) suggests that the burden of XDR *P. aeruginosa* infections in the Philippines may be largely driven by a local lineage of the international epidemic clone ST235. A recent study in a hospital in Jakarta, Indonesia analysed the population composition of *P. aeruginosa* before and after a multi-faceted infection control intervention. While ST235 was the dominant ST pre-intervention, its relative abundance was almost halved in the ten months post-intervention. ^37^ This highlights the importance of hospital infection control and of preventive measures in the use of hospital equipment to contain the further spread of this high-risk clone.

## Acknowledgements

Funding provided by Newton Fund, Medical Research Council (UK), Philippine Council for Health Research and Development.

